# Enrichments along gradients resolve eco-evolutionary forces on subsurface microbiomes

**DOI:** 10.64898/2026.06.29.735320

**Authors:** Zachary S. Cooper, Mingfei Chen, Tiffany Zhao, Jacob J. Valenzuela, Kristopher A. Hunt, Jennifer V. Kuehl, Kathleen S. Walker, Dominique C. Joyner, Daliang Ning, Jizhong Zhou, Terry C. Hazen, Adam P. Arkin, Romy Chakraborty, Nitin S. Baliga

## Abstract

How a single gram of soil harbors billions of microorganisms, each with distinct genomic variants that collectively maintain coherent ecological function(s), is one of microbiology’s grand unsolved problems. A key obstacle is determining which variants contribute to individual- and community-level fitness, in which contexts, and how co-occurring ecotypes interact to divide niche space. Here, using nitrate (NO_3_^-^)-contaminated subsurface sediment as inoculum, we have performed high throughput enrichments in laboratory media of defined carbon source compositions across ecologically relevant gradients of pH and NO_3_^-^. Long-read metagenomics and link-community decomposition of co-occurrence networks of taxa across these enrichments has revealed context-specific functional interactions among dominant generalist and lower-abundance specialist denitrifier ecotypes that comprise 53 distinct enriched communities (EnComs) across 288 enrichments derived from a single sediment sample. We identified a single enzymatic difference of alternative NO_3_^-^ reductases (NapAB vs. NarGHI) with differing substrate affinities that provided a mechanistic explanation for competitive niche partitioning between the two dominant taxa, *Neorhizobium* spp. and *Allorhizobium* spp., along the NO_3_^-^ gradient. Genome-wide polymorphism ratios (pN/pS) revealed that selective pressures vary systematically with carbon source availability and gradients of pH and NO_3_^-^, which helps explain the natural biodiversity and functional interactions of ecotypes within denitrifying communities in the subsurface sediment. Our findings show that controlled enrichments along ecological gradients can thus uncover eco-evolutionary forces of selection, drift, and diversification that sculpt the biodiversity of microbial populations in the natural environment.

## INTRODUCTION

The subsurface environment is inhabited by taxonomically diverse and biogeographically heterogeneous microbial communities, and it is characterized by a dynamic set of multiaxial geochemical gradients that alter the metabolic potential key to global biogeochemical cycling. Oxygen (O_2_) is abundant at the surface-atmosphere interface but is depleted in the shallow subsurface by aerobic respiration [1]. In the deeper O_2_-depleted subsurface, alternative redox compounds, such as nitrate (NO_3_^-^), are used for anaerobic respiration in succession of energetic potential [2]. This vertical stratification of metabolic niche space is the result of a combination of biotic and abiotic forces shaping the physicochemical landscape of the subsurface [3, 4]. In much of the world, anthropogenic inputs to the subsurface, i.e., ammonia (NH_3_) and NO_3_^-^ from agricultural runoff, alter the function of subsurface ecosystems and can result in the increased release of greenhouse gases, such as nitrous oxide (N_2_O), which can have significant effects on the climate, such as ozone depletion and insulating the atmosphere ∼273-310 times more than carbon dioxide (CO_2_) [5, 6]. It is essential to understand the ecological interactions and evolutionary dynamics of microbial communities in response to variable conditions in order to develop a predictive model of subsurface biogeochemical cycling that can be usefully employed for improving environmental management and bioengineering applications [7–13]. Beyond applied biogeochemistry, the mechanisms underlying the eco-evolutionary dynamics of complex microbial communities are fundamental to microbiology.

This study is part of a Department of Energy collaborative effort to deconvolute the complexities of subsurface microbial roles in biogeochemistry, entitled the Ecosystems & Networks Integrated with Genes & Molecular Assemblies (ENIGMA) project, which conducts extensive research on the influence of abiotic and biotic interactions on subsurface microbial communities [14, 15]. Studies in this project have focused on understanding the dynamics of microorganisms in the contaminated subsurface at the ENIGMA Field Research Center (eFRC) in the Y-12 National Security Complex in Oak Ridge, Tennessee, which historically has been the site of many major scientific advancements, including the enrichment of nuclear materials [16]. However, the efforts underpinning these advancements have unintentionally led to the contamination of the local groundwater reservoir with low pH, high NO_3_^-^, and heavy metal-bearing wastewater. More recently, the high concentration of NO_3_^-^ has been found to correlate strongly with the release of gaseous N_2_O to the atmosphere, particularly when pH is low [13, 17–20], making this an ideal test site for investigating the globally-relevant effects of NO_3_^-^ contamination. By conducting geochemical surveys and building hydrogeological models [15, 21, 22], identifying *in situ* microbial community compositions [3, 14, 23], isolating and characterizing endemic microorganisms [24–28], and building synthetic microbial communities (SynComs) for experimental characterization of interactions [4, 20, 29, 30], the ENIGMA project has made tremendous progress in building a fundamental framework for understanding the microbial processes in terrestrial subsurface nitrogen cycling. Yet, there is much that remains to be understood to develop a comprehensive predictive model of microbial function in the dynamic environment of the subsurface [15].

As a respiratory process, denitrification reduces NO_3_^-^ stepwise to nitrite (NO_2_^-^), nitric oxide (NO), N_2_O, and then finally to dinitrogen (N_2_), thus removing a potential environmental contaminant by conversion to a generally neutral atmospheric gas. Each step of this process is genetically encoded by a distinct set of genes, and many microorganisms only encode some of these genes, rendering them unable to complete the full pathway on their own [20, 31]. Given the common occurrence of partial denitrification by individual microbes, cooperative interactions between community members can be essential for the complete conversion of NO_3_^-^ to N_2_, yet the release of N_2_O is commonly observed in the environment where NO_3_^-^ is abundant [13, 19]. Additionally, alternate N-cycle metabolisms, e.g., dissimilatory NO_3_^-^ reduction to ammonium (DNRA) and nitrification, can compete with denitrification by changing the balance of intermediates at the community level, which may result in the release of N_2_O [11, 32]. Given the complexity of the nitrogen cycle and distribution of metabolic potential, it is essential to understand the metabolisms that operate in concert in a taxonomically and genotypically diverse community to develop a complete view of the factors that influence the fate of nitrogen in the environment.

Here, we have applied an enrichment strategy to uncover dynamic context-specific interactions within enriched communities (EnComs) of co-occurring ecotypes across ecologically-relevant gradients of pH and NO_3_^-^. Specifically, using subsurface sediment from the eFRC as inoculum in media with replete or limiting nutrients of simple to complex compositions, we have performed high-throughput enrichments [33–35] across ecologically-relevant gradients of pH and NO_3_^-^, followed by up to three serial transfers interspersed with bottlenecking. Using long-read metagenomics and link-community analysis, we have uncovered how ecotypes of specific taxa with complementary metabolic potential divide niche space by assembling into context-specific EnComs. Further, by quantifying the ratio of nonsynonymous to synonymous polymorphisms (pN/pS) [36, 37], we have characterized how carbon source availability, and gradients of pH and NO_3_^-^ levels have shaped community structure and function at the eFRC by acting combinatorially as eco-evolutionary forces on selection, neutral drift and evolutionary diversification of ecotypes at a genomic-scale, at pathway and gene levels, and all the way down to individual codons. Our findings demonstrate how systematic enrichments with serial transfers and bottlenecking can help uncover how biogeochemical gradients act as eco-evolutionary forces to shape the dynamics of functional interactions among generalized and specialized ecotypes within a heterogeneous and diverse microbial population in the natural environment.

## RESULTS

### Enrichments across eFRC-relevant environmental gradients generate EnComs of overlapping but distinct compositions from the same sediment sample

To dissect microbial interactions that vary across pH and NO_3_^-^ gradients at the eFRC, we simulated field-relevant gradients in multi-well plates to enrich microbial populations using sediment from the vadose zone of the subsurface observatory (SSO; **Fig. 1A**). Three grams of sediment were resuspended in 15 mL of 30 mM phosphate buffer (amended with 5 mM pyrophosphate buffer) and sonicated for 30 minutes in an iced water bath to detach microbes. A 40 μL aliquot of the suspension was used to inoculate each well of 48-well culture plates in an anaerobic chamber, containing 360 µL of either the Northen Lab Defined Medium (NLDM) [38] or Synthetic Groundwater (SGW), both of which were specifically formulated to study subsurface microbiology [39, 40]. Enrichments in NLDM as the base medium included a defined suite of 64 carbon sources at low concentrations (1.8 – 299.5 mg/L) [38], while enrichments in SGW media were amended with single carbon sources (glucose, acetate, citrate, lactate, or pyruvate; **Supplementary Table 1**). Further, enrichments were performed at three different pH levels (4.5, 6, or 7) and over a gradient of NO_3_^-^ concentrations (0, 5, 10, 15, 20, 30, or 40 mM) to simulate environmental transects at the eFRC, which are more extreme (pH: 2.72 – 10.1; NO_3_^-^: ∼ 0 – 139 mM) relative to the SSO (pH: 5.3 – 6.9; NO_3_^-^ : 0.03 – 3 mM; **Fig. 1B**) from where the sediment was procured. The culture plates were incubated anaerobically at 30 °C for 7 days, and a 40 µL aliquot of the culture from each well was transferred (10% inoculum) to the same condition in a new 48-well plate. Altogether, three serial transfers were performed, bottlenecking the community at each transfer point to generate EnComs with members selected by the specific pH, NO_3_^-^ and carbon source conditions within each well (**Fig. 1B**).

**Figure 1.**
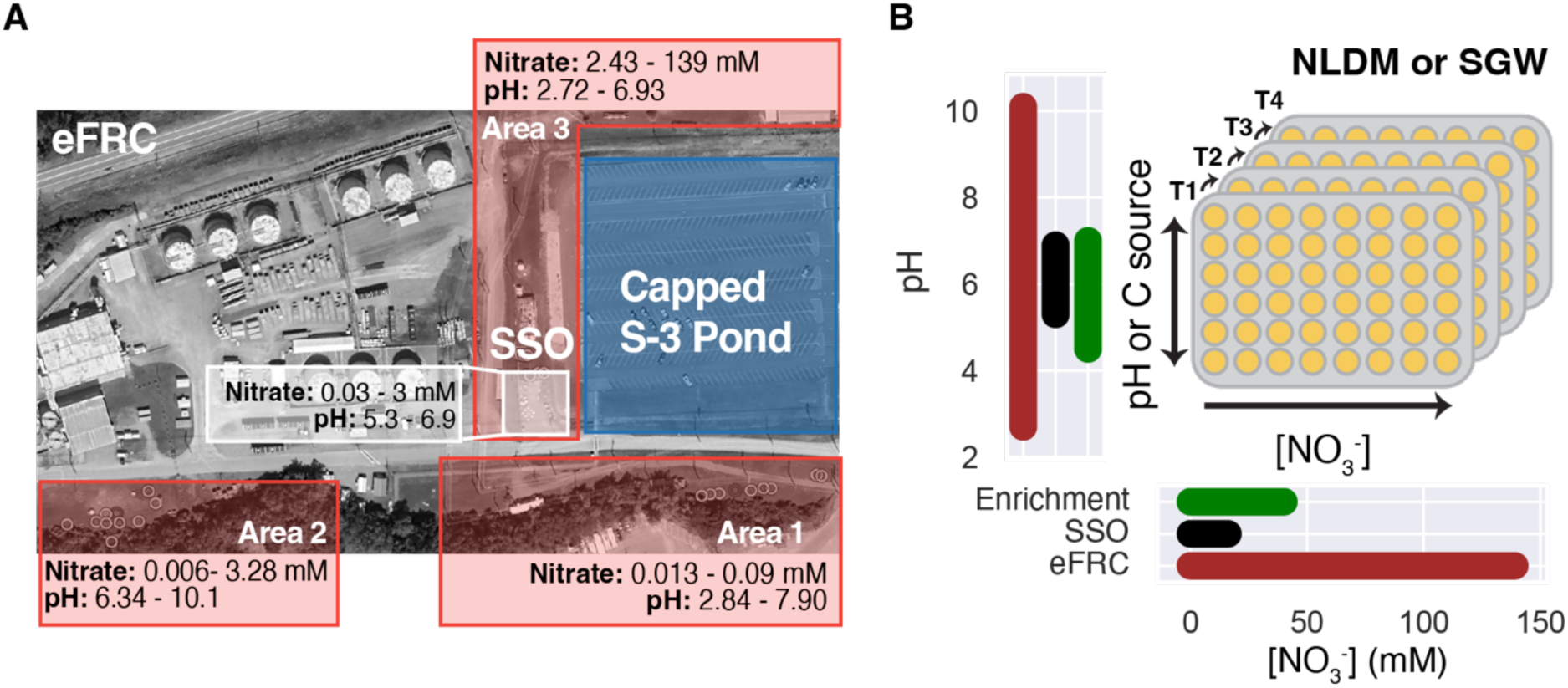
**A**. Aerial view of the eFRC. Range of [NO_3_^-^] and pH across different areas of the eFRC is shown in the inset. The SSO is comprised of 9-wells arranged in a 3×3 grid (1-9), with three wells each in the upper (U), middle (M), and lower (L) rows. The sediment sample used as inoculum in this study was obtained from the vadose zone of the M6 well. **B**. 48-well plate layout using either NLDM or SGW, including gradients of pH, NO_3_^-^ and C-sources (for SGW), and serial-transfer schema for generating EnComs using M6 sediment inoculum

We conducted shotgun metagenomics using long-read sequencing to determine the taxonomic and genomic composition of 288 enrichments (**Supplementary Table 2**; https://narrative.kbase.us/narrative/262281). Co-assembled metagenomes were divided into 41 bins based on hierarchical clustering of gene coverage across samples and k-mer frequencies and were manually refined using Anvi’o (**Supplementary Table 3**; bin annotations and summaries available at https://github.com/baliga-lab/sediment-encoms) [41]. Altogether 23 of the 41 bins represented metagenome-assembled genomes (MAGs; here defined as a bin that represented a distinct population-level genome calculated to be >50% complete using CheckM; **Fig. 2A**; https://narrative.kbase.us/narrative/262281). Using the mean read coverage for a bin in each sample, we calculated the relative abundances of taxa represented in the enrichments across the range of pH, carbon source, and NO_3_^-^ conditions (**Fig. 2B**). Based on homology to universal marker genes annotated in the Genome Taxonomy Database (GTDB), most genera observed across all enrichments were consistent with 16S rRNA amplicon sequence variant (ASV) based assessment of biodiversity at the eFRC [26]. Though, the corresponding ASVs were detected at low average relative abundances in the sediment closest to the inoculum (up to ∼1.44%). This was especially true for the most abundant bin across enrichments, which was identified as *Neorhizobium petrolearium* (100% 16S rRNA identity) and accounted for an average relative abundance of ∼65% across all samples [42]. ASVs corresponding to *Neorhizobium* spp. were detected in varying abundance at many locations across the eFRC (∼0.60% average and ∼1.96% maximum), and in the SSO M6 well sediment they were detected at a relative abundance of ∼1.44%, the highest *in situ* relative abundance of the enriched genera.

**Figure 2.**
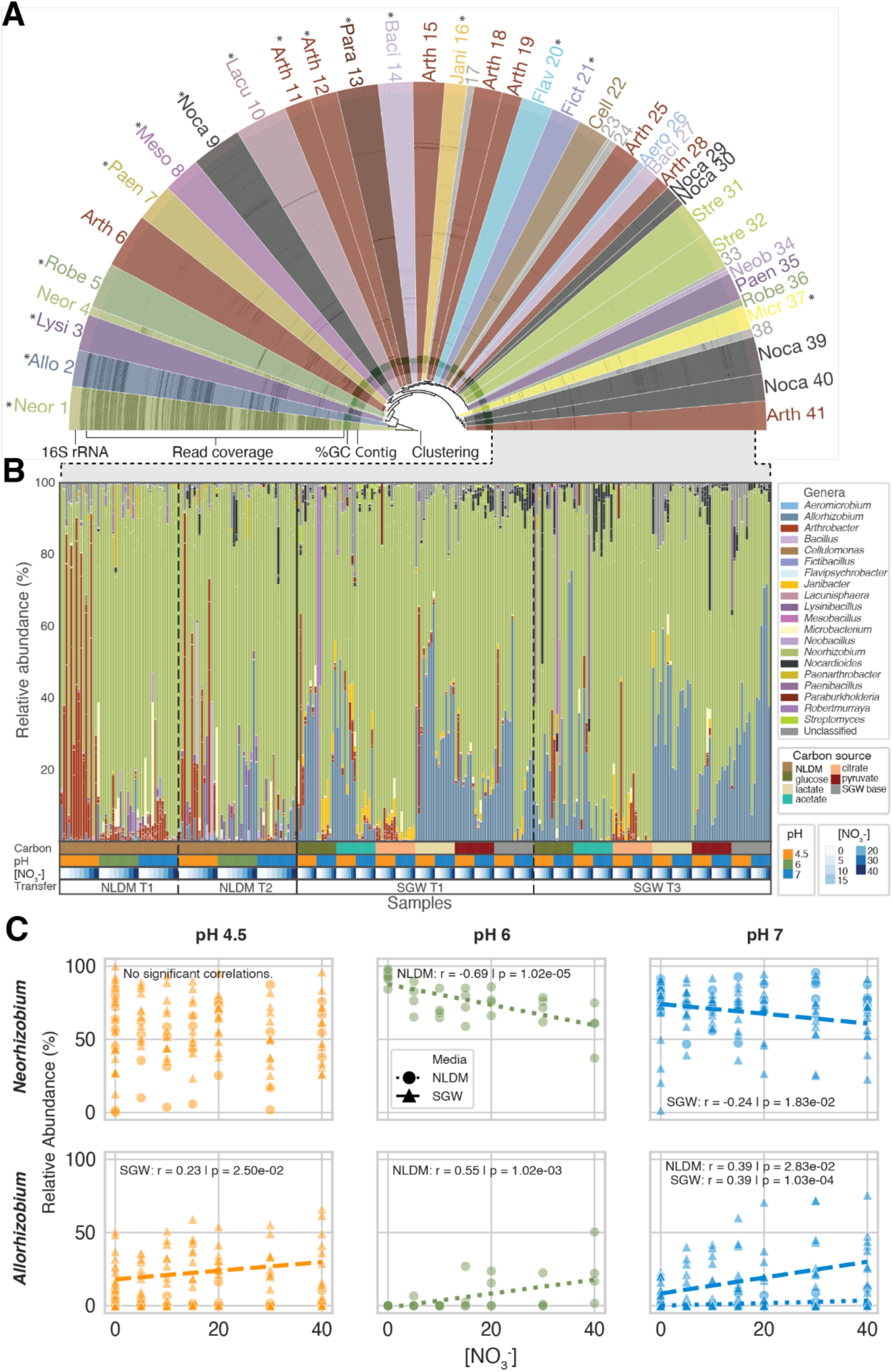
**A**. Metagenomic mapping, binning, and relative abundance of genera in EnComs. Sectors represent bins (labeled by genus -see inset key for details) based on hierarchical clustering of read coverage and k-mer frequencies. Circular tracks indicate relative 16S rRNA gene detection, read coverage in each sample (see panel D for zoomed in view), GC composition, and contigs. **B**. Stacked bar plot illustrates the relative abundance of genera in each sample. Samples are organized by enrichment context: carbon source, pH, NO_3_^-^, base growth medium (NLDM or SGW), and time of sampling vis-a-vis serial transfer -see inset color-coded key for details. **C**. Relative abundance of *Neorhizobium* spp. and *Allorhizobium* spp. as a function of NO_3_^-^ levels and pH (colors corresponding to pH key in panel B).

Relative compositions of multiple genera differentiated EnComs across NLDM and SGW. Whereas *Arthrobacter* spp., *Bacillus* spp., *Lysinibacillus* spp., *Mesobacillus* spp., and *Cellulomonas* spp. were more abundant in NLDM (p < 0.05), *Janibacter* spp., *Streptomyces* spp., and *Nocardiodes* spp. were more abundant in SGW (p < 0.05). Notably, *Allorhizobium* spp., the second most abundant genus across all samples (∼14.1%), was enriched in significantly higher proportion within enrichments in SGW base media than in NLDM (19.5% vs. 3.23%; p = 2.3 x 10^-52^, t-test). Within SGW, the relative abundances of genera were distinct across the different carbon sources. Abundance of *Neorhizobium* spp. was highest with citrate (90.5%), and lowest with lactate (50.8%) relative to all other carbon sources (p < 0.05) (**Supplementary Table 5**). In contrast, abundance of *Allorhizobium* spp. was highest in SGW with lactate (38.4%) and lowest with citrate (0.13%).

We further investigated correlations and co-occurrence patterns for insight into the contexts in which the taxa may assemble and functionally interact as communities. The relative abundance of taxa varied significantly across the three pH conditions (**Fig. 2B**). For example, *Neorhizobium* spp. were enriched in higher relative proportion in pH 6 and 7 than 4.5 (p < 0.05), while *Allorhizobium* spp. and *Arthrobacter* spp. were more abundant at pH 4.5 than 6 or 7 (p < 0.05). Across all samples, the abundances of *Neorhizobium* spp. were significantly negatively correlated with the concentration of NO_3_^-^ (r = -0.14; p = 1.5 x 10^-2^), while *Allorhizobium* spp. were positively correlated with NO_3_^-^ (r = 0.26; p = 5.63 x 10^-6^). The consequence of NO_3_^-^ level changes on composition of enrichments was even more apparent in specific pH and media contexts (**Fig. 2C**). Relative abundance changes in *Neorhizobium* spp., for instance, were significantly negatively correlated with NO_3_^-^ at pH 6 in NLDM (r = -0.69; p = 1.02 × 10^−5^) and pH 7 across SGW media (r = -0.24; p = 1.83 × 10^-2^), but was uncorrelated at pH 4.5 (**Fig. 2C**). Conversely, *Allorhizobium* spp. were significant only at pH 6 (r = 0.55; p = 1.02 × 10^−3^) and 7 (r = 0.39; p = 2.83 × 10^-2^) in NLDM, and the correlations across SGW base media were significant at both pH 4.5 (r = 0.23; p = 2.5 × 10^-2^) and 7 (r = 0.39; p = 1.03 × 10^-4^). None of the other bins were significantly correlated with NO_3_^-^. These findings indicated that while NO_3_^-^ availability was the major determinant of abundances of the dominant genera, pH and carbon sources played more nuanced roles in shaping the overall community structure.

In sum, these findings suggested that the enrichment patterns along biogeochemical gradients may provide clues into how niche space is likely divided to support co-existence of multiple taxa, some of which have functionally redundant genome-encoded capabilities. In subsequent sections, we explore co-occurrence patterns of these taxa across gradients to elucidate which ones likely engage in context-specific, functional interactions as microbial communities.

### Context-specific enrichment and co-occurrence patterns suggest ecologically-distinct functional interactions within EnComs

We aimed to uncover functional interactions among taxa to identify enriched communities (**EnComs**) within enrichments and, more importantly, uncover how these interactions varied across ecologically-relevant chemical transects. We constructed correlation networks using the Fastspar implementation of the SparCC algorithm in a bootstrapping approach to uncover significant correlations in metagenomic coverage of MAGs [43, 44] across samples, while accounting for compositional bias of sparsely correlated taxa [45, 46] (**Fig. 3A**). The edges of the resulting networks were then decomposed using link-community analysis to discover context-specific EnComs based on shared links among interacting populations [47].

**Figure 3.**
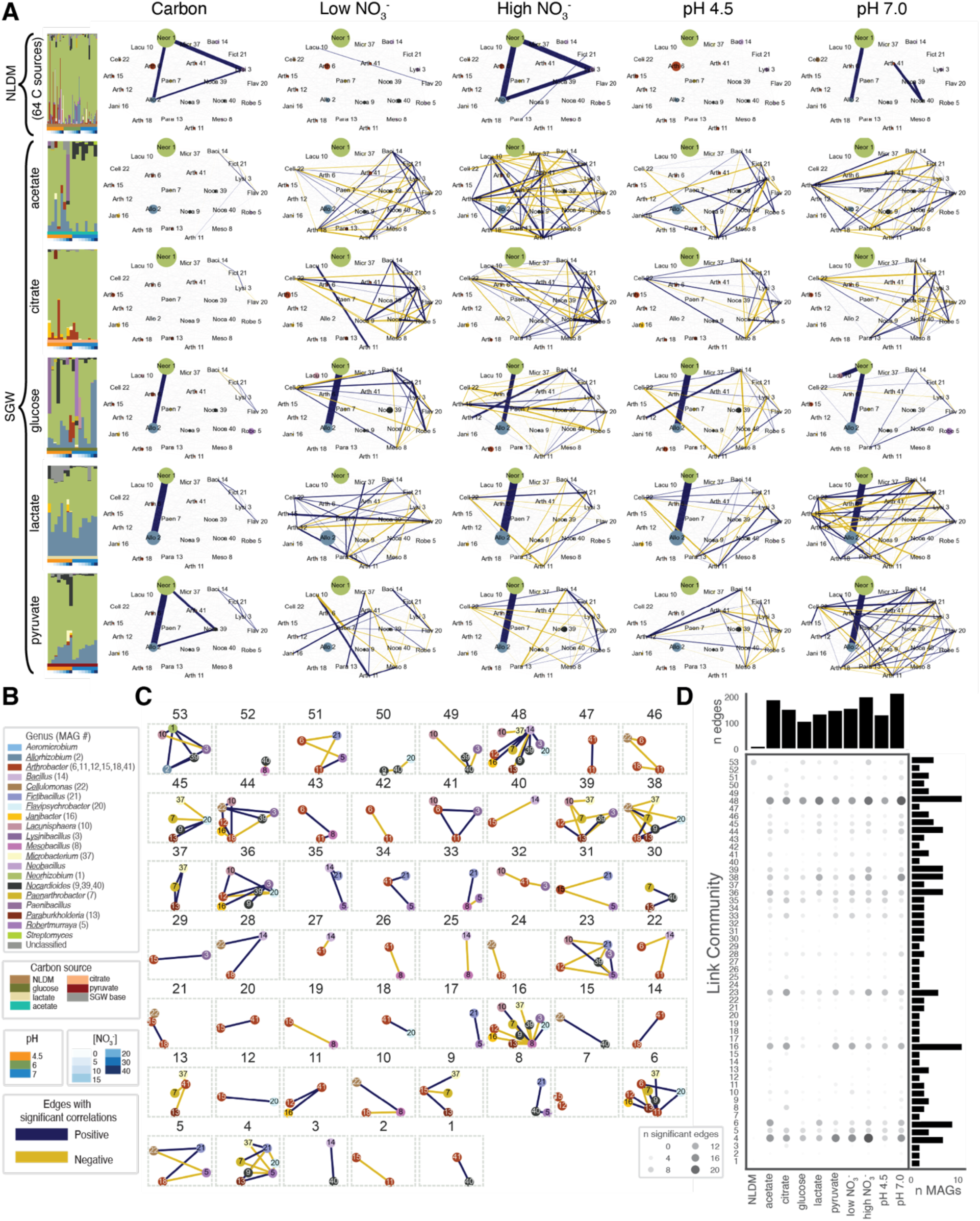
Discovery of EnComs comprised of taxa with similar context-specific co-occurrence patterns and associations discovered with link-community analysis. **A.** Matrix of conditions and corresponding correlation (SparCC) networks; each row represents a single C source and each column represents a subset of conditions. Bar plots (reproduced from Fig 1) indicate relative abundance of genera across conditions represented in the corresponding row. Network edge colors indicate positive (blue) and negative (yellow) correlations (gray edges are not significant), edge thickness is proportional to the correlation coefficient, and node size is proportional to the average relative abundance of that MAG in that subset of conditions. **B.** Key for interpreting color codes for genera (bar plots in panel A, and node colors in panels A and C); MAG number(s) associated with each genus (node labels in panel A: abbreviated four-letter taxon and MAG number, node labels in panel C: MAG number); enrichment media conditions in bar plots, and color code for edges). **C.** Composition and association architecture of taxa within 53 EnComs discovered by link-community analysis. The number on top of each sub-panel is the EnCom ID. Within each EnCom association graph, the node color denotes genus ID and numbers within nodes indicate specific MAG associated with the taxon. **D.** UpSet plot of the number of significant edges (size and shade) among taxa for EnComs in each enrichment context. The right bar plot represents the number of MAGs captured in each link community. The top bar plot shows the number of edges that are significant for each condition.

First, we investigated context-specific co-occurrence patterns among the 23 MAGs using samples from later stages of enrichment (timepoints T2 in NLDM and T3 in SGW), split into 32 distinct conditions based on carbon source, pH (4.5, 6 and 7) and NO_3_^-^ concentrations (low: 5, 10, 15 mM; and high: 20, 30, 40 mM). The analysis discovered up to 69 significant correlations (out of 253 possible pair-wise assessments) in each context (**Fig. 3A**). Although on average a larger number of distinct populations were enriched in NLDM (∼12.7 MAGs detected per condition) relative to SGW with single carbon sources (∼7.1 – 10 MAGs; t-test p < 0.005), in general, fewer significant correlations among MAGs were observed in NLDM (n = 0 – 4), compared to SGW (n = 6 – 69; p < 0.05). The strong correlations between MAGs associated with *Neorhizobium* spp. (Neor 1) and *Allorhizobium* spp. (Allo 2) across most conditions, was consistent with widespread distribution of these genera across many environments, including hydrocarbon contaminated soils [48, 49] and agricultural soils [50–52], where they are considered biomarkers of a healthy rhizosphere [53]. However, we also observed striking differences in correlation patterns across contexts. Notably, Neor 1 and Allo 2 were observed to have significant three-way correlations with different populations depending on the enrichment context. For example, the associations between Neor 1, Allo 2, and *Lysinibacillus* spp. (Lysi 3) were significant across all NLDM samples, but detected neither at low NO_3_^-^ nor across all SGW media conditions with single carbon substrates. Instead, in SGW media enrichments, Neor 1 and Allo 2 were significantly correlated with *Nocardioides* spp. (Noca 39) across all contexts when pyruvate was the carbon source, and with *Lacunisphaera* spp. (Lacu 10) with glucose as the carbon source only in pH 7 conditions.

Next, we calculated mean pairwise correlation values between MAGs and the number of conditions in which each pair was significantly correlated (**Fig. 3A**). Correlation analysis of single-carbon-source enrichments, stratified by NO_3_^-^ and pH, revealed significantly more positive and negative inter-population correlations than observed overall (p = 1.8 × 10^-4^). Notably, the number of correlations were highest in enrichments at high NO_3_^-^ and at neutral pH (n = 17 – 69), relative to enrichments across all conditions with the same carbon substrate (n = 6 – 10). We posit that the strong context-dependence of inter-population correlations in specific pH and NO_3_^-^ conditions is likely indicative of functional microbial assemblages occupying corresponding ecological niches in the subsurface at the eFRC.

Subsequently, we used link-community clustering [47] and identified MAGs with similar co-occurrence patterns to delineate EnComs and identify the specific contexts in which the included members likely interacted. Using this framework, we identified 53 EnComs with varying population memberships across contexts (**Fig. 3C-D**), revealing a structural hierarchy of potential interactions between dominant and lower-abundance populations. Neor 1 and Allo 2 clustered together into a single EnCom (#53), which also included lower-abundance populations (Lysi 3, Lacu 10, and Noca 39) whose relative abundances were strongly correlated with the dominant populations in specific subsets of conditions. This pattern suggests primary cooperative and competitive dynamics between Neor 1 and Allo 2 across the NO_3_^-^ gradient, with more commensal relationships with the lower-abundance co-occurring populations.

In NLDM, which includes 64 distinct carbon sources, EnComs were defined largely by strong positive correlations among the two dominant populations, with relatively few significant associations involving lower-abundance populations. In contrast, single-carbon-source media yielded more complex EnCom structures characterized by numerous significant associations among non-dominant populations, suggesting greater metabolic interdependence under resource-limited conditions. Consistent with this complexity, lower-abundance populations were distributed across multiple EnComs of 2–10 members each, with membership patterns defined primarily by co-occurrence in single-carbon media at pH 7 and high NO_3_^-^ concentrations (**Fig. 3D**). For example, EnCom #48 is centered on Baci 14, which correlates positively with Noca 39, Jani 16, and Micr 37, and negatively with Lacu 10. Baci 14 is most abundant in NLDM, suggesting it may depend on metabolic by-products from Noca 39, Jani 16, and Micr 37, populations that dominate under single-carbon resource-limited conditions, while competing with Lacu 10 for shared metabolites. Similarly, EnCom #4 features a more complex network, with potential cooperative interactions among Micr 37, Fict 21, and Paen 7 balanced against possibly competitive interactions with Noca 9 and Robe 5. The metabolic potential underlying these interaction patterns is explored further in the following section.

Overall, the correlation and link-community analysis revealed a dynamic landscape of probable context-dependent EnComs with likely functional interactions in specific contexts. The disruption of the correlation between *Neorhizobium* spp. and *Allorhizobium* spp. across pH, NO_3_^-^, and carbon conditions (**Fig. 3A**) suggested that while these genera were present in high abundance across most samples, they may have competing roles across specific environmental contexts. This hypothesis is consistent with the context-specific co-occurrence of these dominant genera with other lower abundance genera that also varied across pH and NO_3_^-^ gradients, and by carbon source. While the core community structure remained relatively stable, the specific correlation networks demonstrated remarkable plasticity in response to environmental conditions. Together these findings demonstrate how high-throughput enrichment along controlled pH and NO_3_^-^ gradients with defined carbon sources discovered significant context-specific associations among naturally-co-occurring populations in groundwater microbial communities at the eFRC.

### MAGs of taxa within EnComs encode complementary metabolic capabilities that align with the context of enrichments

We assessed metabolic pathway completeness and gene content of each of 16 MAGs (>90% complete) using KEGG KofamScan orthologs (KOs) and NCBI COG annotations to uncover evidence supporting context-dependent metabolic interactions that underpin link-community uncovered architecture of associations among taxa within EnComs (**Fig. 4A**) [54, 55]. Many of the genera that were detected, including *Neorhizobium* spp., *Allorhizobium* spp., *Arthrobacter* spp., *Bacillus* spp., *Robertmurraya* spp., *Janibacter* spp., *Microbacterium* spp., and *Nocardioides* spp., are broadly characterized by flexible heterotrophic metabolisms [56–63], which was consistent with the observation that nearly all MAGs associated with these taxa encoded complete central carbon metabolism pathways, including glycolysis, gluconeogenesis, the citrate cycle, and the pentose phosphate pathway (**Fig. 4A**; **Supplementary Table 6**). The glyoxylate cycle, a pathway essential for acetate catabolism, was encoded in MAGs of *Neorhizobium* spp., *Allorhizobium* spp., and *Nocardioides* spp. that were abundant in acetate-amended media as well as in media supplemented with other carbon sources, further illustrating the metabolic flexibility of these populations. However, not all EnComs included metabolic generalists. Several genera that were members of multiple EnComs under specific carbon source conditions are known to possess correspondingly specialized metabolisms.

**Figure 4.**
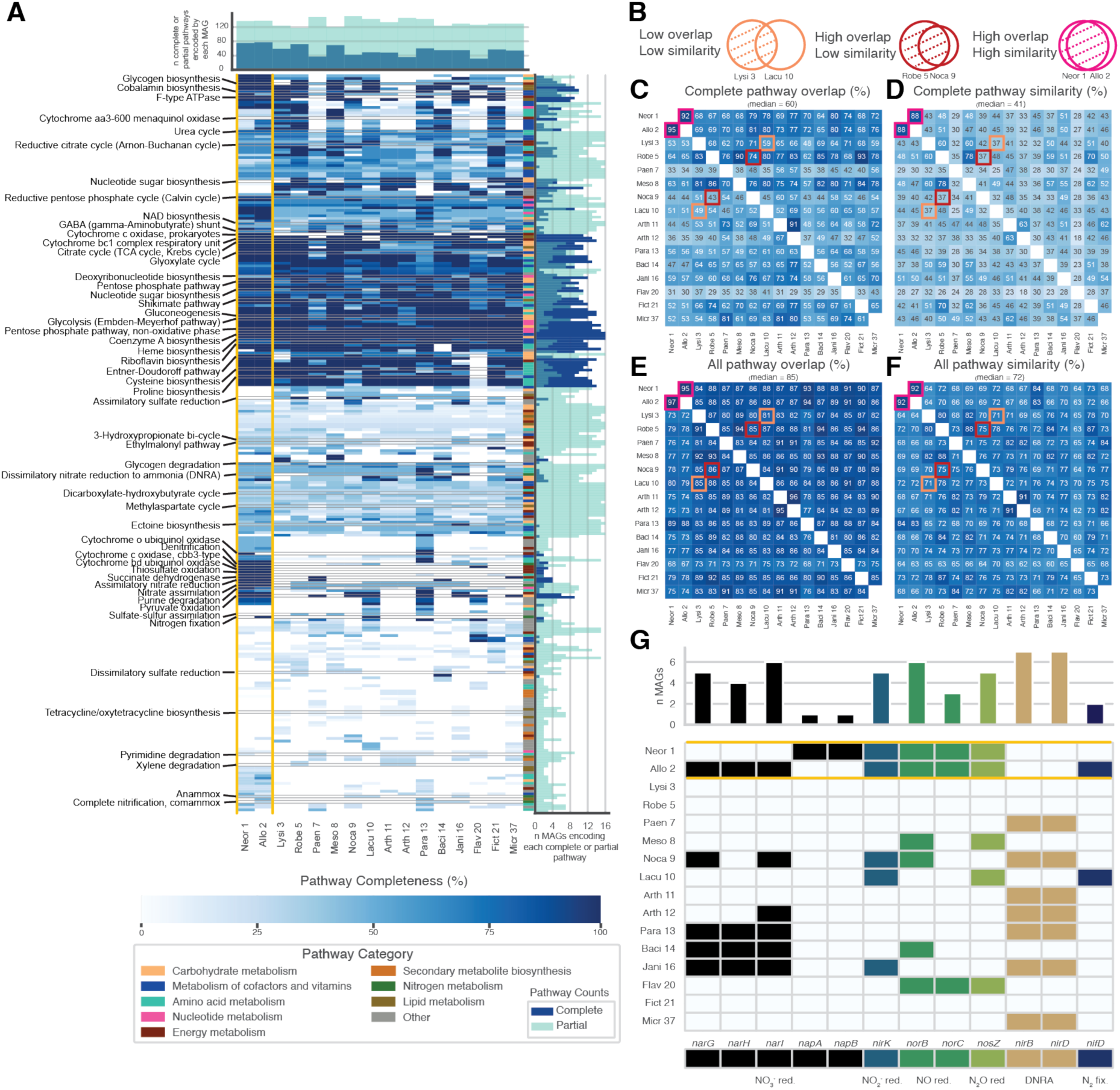
Metabolic pathways encoded across enriched populations with >90% complete MAGs. **A.** Heatmap shows completeness of KEGG metabolic pathways encoded by each MAG. Pathways encoded in Neor 1 and Allo 2 are shown between the two gold vertical borders. Selected pathways with complete sets of genes, and a few incomplete examples, across Neor 1 and Allo 2 are labelled. The bar chart on top shows the number of complete and partial pathways encoded by each MAG, and the chart on the right shows the number of MAGs encoding each pathway. **B.** Venn diagrams illustrate three scenarios of overlap and similarity between pathways encoded by two MAGs with provided examples. “Overlap” is the intersection divided by contents of the left circle, and “Similarity” is the intersection divided by the union of both circles (Methods). Heatmaps show degree of *metabolic pathway overlap*, i.e., the proportions of complete (**C**) and all (partial and complete) pathways (**E**) encoded by MAGs in columns, that are also encoded by MAGs in rows. Heatmaps in **D** and **F**, show *metabolic pathway similarity*, i.e., the percentage overlap of genes in complete and all (partial and complete) pathways encoded by MAGs in columns vs. rows, respectively. **G.** Genes of nitrogen metabolism across MAGs. Bar plot in top panel shows number of MAGs with each gene of denitrification, DNRA and BNF pathways. The heatmap in the bottom panel indicates presence and absence of denitrification, DNRA and BNF genes in each MAG. Nitrogen metabolism genes encoded in Neor 1 and Allo 2 are in the top two rows between the two gold horizontal borders.

Next, we calculated two metrics to evaluate metabolic redundancy (“pathway overlap”) and complementarity (“pathway similarity”) across the high-quality MAGs (**Fig. 4B**, Methods). We investigated if the combination of these two metrics would explain enrichment of MAGs in specific contexts as well as their membership in EnComs. For instance, *Neorhizobium* spp. and *Allorhizobium* spp. MAGs encoded a high proportion of the metabolic pathways found across all other populations, yet they also possessed numerous pathways that were unique, reflecting both broad metabolic overlap with the community and greater similarity to each other than to any other population (**Fig. 4C–F**). This generalist metabolic capacity is consistent with the sustained dominance of *Neorhizobium* spp. and *Allorhizobium* spp. MAGs across most carbon sources and supports the opportunistic utilization of secondary metabolites exuded by the dominant populations to the benefit of lower-abundance community members. The pairwise evaluation of complete metabolic pathways encoded by MAGs within EnComs showed low overlap and low similarity (less than the median value) for some MAGs (e.g., Lacu 10 and Lysi 3 in EnCom #53; **Fig. 4C-F**). Consider, for instance, EnCom #53, which is comprised of populations associated with *Neorhizobium* spp. (Neor 1), *Allorizobium* spp. (Lacu 10), *Lysinibacillus* spp. (Lysi 3) and *Lacunisphaera* spp. (Lacu 10). The reported inability of *Lysinibacillus* spp. to metabolize carbohydrates [64, 65], may explain its enrichment exclusively in NLDM, a medium with diverse nutritional sources [38]. In contrast, *Lacunisphaera* spp. are sugar utilizers [66], consistent with the enrichment of this genus primarily in glucose containing media, where its abundance was significantly correlated with *Neorhizobium* spp. (**Fig. 3A**). The negative correlation in occurrence patterns of Lacu 10 and Lysi 3, and their low metabolic overlap and similarity with each other suggest that these populations have mutually exclusive roles with the two dominant genera. Additionally, we observe MAGs with high overlap but low similarity (e.g., Robe 5 and Noca 9 in EnCom #4; **Fig. 4C-F**) which indicates an imbalance in competitive potential. While both *Robertmurraya* spp. and *Nocardiodes* spp. are broadly generalists, the more expansive repertoire of complete pathways held by Robe 5 provides a higher potential for competition while Noca 9 has more incomplete pathways (**Fig. 4A**) and may more heavily rely on cooperative metabolite exchange. The overlap and similarity values for all pairs of MAGs increase when both complete and partially encoded metabolic pathways are considered (**Fig. 4E-F**), suggesting that functions that cannot be completed individually may be enabled by cooperative metabolic exchange.

Among the high-quality MAGs, we identified nitrogen-cycling metabolisms encompassing denitrification, DNRA, and biological nitrogen fixation (BNF), collectively suggesting a complex network of community-level nitrogen-cycling interactions (**Fig. 4G**). The two most abundant MAGs across all samples, Neor 1 and Allo 2, encoded a complete complement of denitrification pathway genes (*narG* or *napA*, *nirK*, *norB*, and *nosZ*). In contrast, six MAGs affiliated with lower-abundance taxa (Meso 8, Noca 9, Lacu 10, Baci 14, Jani 16, and Flav 20) lacked at least one intermediate enzymatic step. Four MAGs (Noca 9, Para 13, Baci 14, and Jani 16) encoded genes for both the NO_3_^-^- and NO_2_^-^ -reduction steps of DNRA (*narG* and *nirB*, respectively), whereas four additional MAGs (Paen 7, Arth 11, Arth 12, and Micr 37) encoded only the NO_2_^-^-reduction gene (*nirB*), representing partial DNRA capacity. Two MAGs (Allo 2 and Lacu 10) encoded the key nitrogenase subunit gene (*nifD*), indicative of BNF potential.

The apparent incompleteness of several metabolic pathways should be interpreted with caution, as non-circularized MAGs are inherently incomplete and may not capture the full coding repertoire of the represented populations. Nevertheless, partially encoded denitrification pathways have been reported as a common feature of subsurface microbial communities [13, 20, 31], consistent with the view that modular, cooperative metabolism operates at the community level rather than within individual populations. Notably, the Lysi 3 and Robe 5 MAGs encoded no respiratory nitrogen-cycling genes, but do encode oxidoreductases and sulfate reductases as well as lactate and alcohol dehydrogenases for fermentation, suggesting that these taxa may depend obligately on alternative electron acceptors for energy conservation. This is a striking observation given that they represent abundant populations strongly correlated with the dominant community members. Additionally, *Bacillus* spp. are known to have fermentative metabolisms, with capability to occupy respiratory niches that are overlooked when focusing on O_2_ and NO_3_^-^ metabolisms. Fermentative metabolisms also benefit from secondary metabolite interactions and may explain the complex interaction networks observed across single carbon source enrichments and as displayed by Baci 14 across EnComs.

The Neor 1 and Allo 2 MAGs encoded 75 and 77 complete metabolic pathways, respectively, of which 88% of the combined total were shared (**Fig. 4A,D**), indicating a high degree of potential niche similarity between these two dominant taxa. Despite this broad metabolic overlap and similarity, differences in gene content may underlie the divergent competitive and cooperative dynamics observed between these MAGs within the EnComs. Although both MAGs encoded a complete denitrification pathway (**Fig. 4G**), they carry alternative NO_3_^-^ reductase operons: Neor 1 encoded the *napAB* operon (periplasmic NO_3_^-^ reductase complex), whereas Allo 2 encoded the *narGHI* operon (membrane-bound, cytoplasmically oriented NO_3_^-^ reductase complex). The NapAB complex has been demonstrated to exhibit higher substrate affinity for NO_3_^-^ than NarGHI [67–69]. Beyond this distinction, both MAGs encode an otherwise identical complement of denitrification genes, NO_2_^-^ reductase (*nirK*), NO reductase (*norBC*), and N_2_O reductase (*nosZ*), suggesting that this single enzymatic difference may have been a key determinant of their anticorrelation along the NO_3_^-^ concentration gradient in the enrichments. Taken together, the extensive overlap in shared metabolic capacity reflects similar overall niche potential between these co-dominant populations, while the targeted differences in gene content offer mechanistic insight into the molecular basis of their competitive partitioning within the EnComs.

### Population genomics reveals eco-evolutionary forces operating at the genomic, pathway, gene, and codon levels

We hypothesized that genomic diversity of MAGs enriched across gradients in pH (low to neutral), NO_3_^-^ (low to high), and distinct carbon sources would reveal evidence of eco-evolutionary forces acting on taxa in their natural environment. We tested this hypothesis by calculating the ratios of nonsynonymous to synonymous polymorphisms (pN/pS) within all codons in the dominant *Neorhizobium* spp. MAG (Neor 1) across enrichment contexts [36, 37]. We analyzed the distribution of pN/pS values at each codon position for evidence of purifying selection (pN/pS < 1, intensifying as they approach 0), neutral drift (pN/pS = 1), or diversification (pN/pS > 1, intensifying as the value increases) [36, 37] across pathways, at the individual gene level, and down to individual codon positions in specific genes (**Fig. 5**).

**Figure 5.**
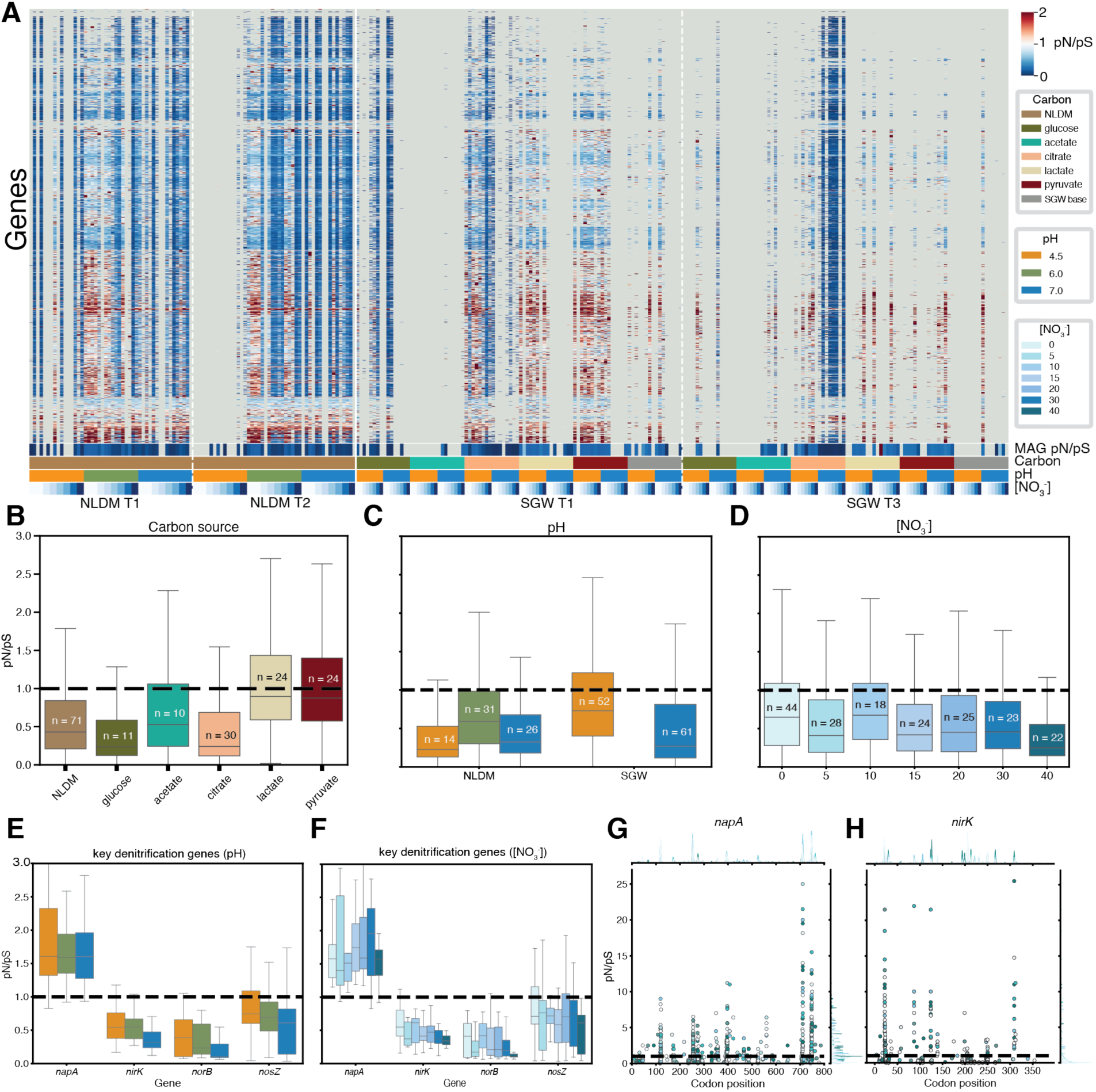
Genomic, gene, and codon-level pN/pS of the Neor 1 MAG across enrichment conditions. **A.** Row-clustered heatmap of pN/pS values for all genes. Tracks at the bottom of the heatmap show mean pN/pS values across all genes, and the context of enrichments vis-à-vis carbon source, pH and NO_3_^-^ levels. **B-D.** Boxplots show distributions of pN/pS values of all genes encoded by Neor 1 in enrichments across carbon sources **(B),** pH **(C),** and NO_3_^-^concentrations **(D).** Each box is labelled with the number of samples with data covered by that condition. **E-F.** Distribution of pN/pS values for each denitrification pathway gene encoded by Neor 1 across enrichments at pH 4.5, 6.0, and 7.0 **(E)** and across NO_3_^-^ concentrations **(F). G-H.** pN/pS values at each codon position within napA **(G)** and nirK **(H)** colored by NO_3_^-^concentrations. The top panel shows the density of pN/pS values at each codon, and the right panel shows the density of pN/pS values across the whole gene.

We observed significant differences in eco-evolutionary forces acting on all genes across the entire MAG, across transfers, between media and carbon sources, at each pH, and along the NO_3_^-^ gradient (p < 1 × 10^−18^; **Fig. 5B-D**). While enrichments in NLDM were generally associated with greater purifying selection (mean pN/pS = 0.69), in SGW supplemented with single carbon sources there was evidence of a diversity of eco-evolutionary forces acting on Neor 1, with mean pN/pS values ranging from 0.59 – 1.36 for each carbon source; **Fig. 5B**). As expected, greater purifying selection was observed in successive transfers, with the mean genomic pN/pS decreasing significantly from 0.72 to 0.67 in NLDM from the first to second transfer (p < 1 × 10^−18^), and from 0.99 to 0.89 in SGW from the first to third transfer. Our findings also uncovered significant differences in eco-evolutionary forces of pH and NO_3_^-^ on Neor 1 across NLDM and SGW (**Fig. 5C-D**). In NLDM, relative to pH 6.0 (pN/pS = 0.83), the purifying selection across genomic loci of Neor 1 was significantly higher across enrichments at pH 4.5 (pN/pS = 0.49) and pH 7 (pN/pS = 0.58) (p < 1 × 10^−21^ for all comparisons). By contrast, we observed evidence of evolutionary diversification of Neor 1 across SGW enrichments at pH 4.5 (pN/pS = 1.11) and purifying selection at pH 7 (pN/pS = 0.73; p < 1 × 10^−100^). Across the NO_3_^-^ gradient, pN/pS declined significantly from 0.93 at the lowest concentration to 0.52 at the highest (r = –0.07; p < 1 × 10^-100^), indicating stronger purifying selection at elevated NO_3_^-^ concentrations. In general, genes with signatures of evolutionary diversification (pN/pS > 1) were pre-dominantly involved in the transport of organic substrates, central carbon metabolism (e.g., glycolysis and the TCA cycle), and amino acid biosynthesis and degradation. By contrast, genes under strongest purifying selection (i.e., lowest overall mean pN/pS values) were associated with core functions, such as transcription and translation, likely because enrichments typically favor variants with maximum fitness (i.e., variants with fast growth).

Next, we inspected eco-evolutionary forces acting specifically on denitrification pathway genes (**Fig. 5E-F**). Across all media and pH contexts, *napA* exhibited a strong signature of evolutionary diversification (pN/pS mean = 1.98), while *nirK*, *norB*, and *nosZ* were under purifying selection with significantly lower pN/pS values (*nirK* = 0.50, *norB* = 0.33, *nosZ* = 0.85, p < 2.7 × 10^-15^). The purifying selection on *nirK*, *norB*, and *nosZ* was greater at pH 7.0 relative to pH 4.5 (p < 0.004; **Fig. 5E**). Interestingly, the purifying selection of *nirK* was greater with increasing NO_3_^-^concentrations (pN/pS values decreasing from 0.59 (0mM NO_3_^-^) to 0.37 (40mM NO_3_^-^); r = –0.29, p = 0.008; **Fig. 5F**), while eco-evolutionary forces on rest of the denitrification pathway genes stayed constant. Given their contrasting eco-evolutionary signatures and their roles in the first two steps of denitrification, we examined codon-level selection in *napA* and *nirK* (**Fig. 5G-H**). Although the codon-level analyses identified sites of evolutionary diversification in both *napA* and *nirK*, no consistent gradient-associated pattern emerged across either pH or NO_3_^-^ concentration, suggesting that selection acts broadly across these genes rather than at environmentally-tuned hotspots.

Although some variants may have arisen during the course of laboratory enrichment, we posit that the majority of observed polymorphisms reflect standing genomic diversity present in the natural eFRC population, and that pre-existing variants conferring greater fitness were selectively enriched in specific chemical contexts. In this way, the pN/pS patterns documented here offer a window into the eco-evolutionary forces that have shaped microbial biodiversity at the eFRC in situ. By systematically evaluating the relative contributions of purifying selection, neutral drift, and diversifying selection across ecologically relevant gradients of pH, nitrate, and carbon source, we reveal how niche space is finely partitioned among ecotypes that have adapted to the distinct selective pressures experienced in the subsurface with pressures faithfully recapitulated, at least in part, by the controlled chemical conditions of enrichments.

## DISCUSSION

### Interpreting microbial biodiversity through eco-evolutionary dynamics in controlled enrichments

Natural microbial communities harbor extraordinary taxonomic and genomic diversity spanning from the kingdom to the codon level, yet translating this diversity into a mechanistic understanding of how specific variants contribute to individual- and community-level fitness remains a fundamental challenge in microbial ecology [3, 14, 23]. A major source of this difficulty is the spatiotemporally complex biogeochemistry of natural environments, where simultaneous variation in diverse factors, including pH, electron acceptor availability, carbon source composition, and nutrient stoichiometry, collectively shapes organismal and community fitness across both short adaptive and long evolutionary timescales [3, 15]. The resulting difficulty of disentangling causality from correlation in field data has motivated the development of controlled experimental platforms that preserve biological complexity while permitting systematic perturbation of individual environmental variables [33–35]. By performing enrichments to uncover EnComs from subsurface sediment in media of defined composition across ecologically relevant chemical gradients, this study demonstrates how variants of diverse, naturally co-occurring microbial species partition niche space at both the individual population and community levels, and how population-level genomic diversity reflects the cumulative product of eco-evolutionary forces including purifying selection, neutral drift, and evolutionary diversification.

### Community assembly reflects the intersection of carbon source complexity and nitrogen redox chemistry

Microbial communities are known to exhibit considerable taxonomic variability while often maintaining relatively stable functional community structures, a phenomenon described as functional redundancy [70]. Our findings both confirm and elaborate on this paradigm. The broad taxonomic structure of the EnComs was dominated by *Neorhizobium* spp. and *Allorhizobium* spp. across most enrichment conditions regardless of pH, NO_3_^-^ concentration, or carbon source, suggesting a conserved functional core anchored by these two populations. Yet the quantitative shifts in the relative abundances of both dominant and subsidiary populations across chemical gradients reveal that this apparent functional conservatism conceals finely partitioned niche space operating at the level of enzymatic variants rather than metabolic categories.

The role of carbon source complexity in structuring the EnComs was particularly striking. In NLDM, which presents a diverse array of carbon substrates at low individual concentrations [38], more taxa achieved significant relative abundances simultaneously, and the network of significant correlations between taxa was comparatively sparse. By contrast, enrichments in SGW supplemented with single carbon sources generated more complex, context-specific correlation networks among the non-dominant populations. This pattern is consistent with the principle that resource diversity reduces competitive exclusion by supporting distinct metabolic niches simultaneously [71, 72], while resource limitation under single-substrate conditions intensifies competitive dynamics and forces remaining taxa into tighter ecological dependencies, including cross-feeding on metabolic exudates generated by dominant populations [73]. The observation that taxa such as *Lacunisphaera* spp., which were characterized as specialized sugar utilizers [66], were enriched specifically in glucose-containing media, while *Lysinibacillus* spp., which lack the capacity to metabolize carbohydrates [64, 65], were essentially restricted to NLDM, provides strong evidence that the communities assembled in these enrichments reflect ecologically real preferences evolved in the natural environment.

The influence of pH on community composition adds a further axis of niche differentiation. *Neorhizobium* spp. were enriched at circumneutral pH (6–7), whereas *Allorhizobium* spp. and *Arthrobacter* spp. were relatively more abundant at pH 4.5. This pH-dependent partitioning of dominant populations is consistent with observations across multiple subsurface and soil environments, where pH is among the strongest predictors of microbial community composition [23]. At the eFRC, acidification of groundwater from historical wastewater inputs has been shown to correlate with increased N_2_O emissions [13, 17–20], and the pH-dependent shift in dominant denitrifier populations documented here has direct implications for the efficiency and completeness of nitrogen removal from contaminated groundwater. The combination of pH and NO_3_^-^ concentration thus co-determines the metabolic landscape of denitrifying communities in ways that can plausibly account for the variable stoichiometry of N_2_O production observed in field measurements at the eFRC [13, 20].

### Enzymatic variants underpin niche partitioning between functionally overlapping co-dominant taxa

One of the central findings of this study is that MAGs associated with *Neorhizobium* spp. (Neor 1) and *Allorhizobium* spp. (Allo 2), despite sharing 88% of their complete metabolic pathways and co-dominating the EnComs across most conditions, exhibit a robust anticorrelation along the NO_3_^-^gradient that can be mechanistically explained by a single enzymatic difference: the use of the periplasmic, high-affinity NapAB nitrate reductase complex by the *Neorhizobium* spp., versus the membrane-bound, lower-affinity NarGHI complex by the *Allorhizobium* spp. [67–69]. Recent work by Crocker et al. [35] showed that pH has a distinct effect on the abundance of these enzymes across global soil metagenomes, where *narG* genes decreased with increasing pH and *napA* genes increased with pH. This is consistent with our findings that *narG*-encoding *Allorhizobium* spp. were more abundant at low pH while *napA*-encoding *Neorhizobium* spp. were more abundant at higher pH. Additionally, Crocker et al. [35] demonstrated a pH-dependent effect on the rate of NO_3_^-^ reduction for isolated bacteria encoding each enzyme, where a *narG*-encoding *Pseudomonas* sp. reduced NO_3_^-^ at a lower initial rate than a *napA*-encoding *Rhizobium* sp.. Our results indicate that the concentration of NO_3_^-^ plays a compounding role in niche partitioning, where *Neorhizobium* spp. abundance declined with increasing NO_3_^-^ at pH 6 and 7, while *Allorhizobium* spp. abundance increased. This pattern can be interpreted as competitive dominance of the high-affinity NapAB system at low substrate concentrations giving way to the NarGHI system when NO_3_^-^ is replete and affinity differences are less consequential [67, 68]. This enzymatic-level mechanism for the coexistence of two otherwise nearly metabolically identical populations is a compelling example of the ecotype concept in action, where sympatric populations are maintained by resource-based niche partitioning operating at very fine genetic resolution [43, 74].

The broader implication of this finding is that functional redundancy at the metabolic pathway level does not imply ecological equivalence. Subtle differences in enzyme kinetics, regulatory architecture, or environmental optima can generate sufficient niche differentiation to sustain coexistence across resource gradients [70]. This principle is well-documented in ocean systems, for example, in the partitioning of *Prochlorococcus* spp. ecotypes along light and nutrient gradients [75], and of *Vibrio* spp. populations along temperature, salinity, and particle-size niches [76], but its demonstration in a controlled enrichment setting with community-derived populations from a contaminated subsurface environment provides direct experimental evidence of its generality. Critically, resolving this ecotype-level niche partitioning required maintaining population-level genetic heterogeneity from the natural environment through the experimental system, a capacity that bottom-up approaches using clonal isolates fundamentally cannot provide [4, 33, 35].

The partial denitrification capacity encoded by many of the lower-abundance MAGs further illustrates how metabolic complementarity structures community-level nitrogen cycling. As demonstrated in prior work at the eFRC and other denitrifying systems [20, 31], the cooperative handoff of nitrogen intermediates among partial denitrifiers reduces the energetic cost of individual populations while enabling complete conversion of NO_3_^-^ to N_2_ at the community scale. The competition between denitrification and DNRA for shared intermediates such as NO_2_^-^ adds an additional ecological dimension, with DNRA-capable populations potentially buffering against toxic intermediate accumulation under high-flux conditions [32, 77]. The community-encoded nitrogen cycle revealed here thus represents a distributed metabolic architecture whose emergent function of bulk nitrogen removal depends not on any single population but on the collective, context-dependent assembly of taxonomically diverse co-occurring ecotypes.

### Genomic signatures of selection reveal the eco-evolutionary forces shaping population diversity

The analysis of genetic variability of *Neorhizobium* spp. populations across enrichment contexts provides insight into a central question in microbial ecology: how does the extraordinary genomic diversity observed in natural environments arise and persist? Our findings support a model in which the natural environment simultaneously maintains diverse selective forces at different genomic loci, such that the biodiversity captured in field metagenomes reflects the cumulative product of purifying selection, neutral drift, and diversifying selection operating in context-dependent manners [36, 37].

At the genome-wide level, enrichments in NLDM, in which diverse carbon sources are available simultaneously, were associated with greater mean purifying selection (lower pN/pS) than single-substrate SGW enrichments, suggesting that nutritional diversity relaxes the selective constraints on genome-wide optimization and permits a greater range of genetic variants to persist. The particularly strong purifying selection observed at pH 4.5 in NLDM, and the evidence for evolutionary diversification at pH 4.5 in SGW, implies that physiological stress, i.e., acidity in this context, interacts with resource availability to modulate the strength and direction of selection across the genome in ways that would be effectively invisible to field sampling alone. The progressive intensification of purifying selection across successive enrichment transfers is consistent with the expectation that serial bottlenecking under defined conditions reduces effective population size and allows selection to act more efficiently on standing variation.

At the gene level, the strikingly high pN/pS of *napA* (mean 1.98) across all conditions, relative to the much lower values for downstream denitrification genes (*nirK*, *norBC*, *nosZ*), reveals a pathway-specific pattern of selective pressure that tracks substrate availability rather than whole-genome physiological state. When NO_3_^-^ is abundant, its reduction at the first step of denitrification is unlikely to be rate-limiting, and variants of *napA* that alter catalytic efficiency carry lower fitness consequences, permitting diversification. In contrast, the downstream intermediates (NO_2_^-^, NO, and N_2_O) are present at lower and more variable concentrations, making enzymatic efficiency at those steps more consequential for fitness and therefore subject to stronger purifying selection [78–80]. This substrate-availability model for differential selective pressure along a metabolic pathway provides a general interpretive principle for understanding genomic diversity in environmentally-derived populations, such that genes whose substrates are rarely limiting should accumulate nonsynonymous variants more freely than genes acting on transiently available or potentially toxic intermediates.

These observations are consistent with a broader eco-evolutionary framework for understanding microbial biodiversity in natural environments. Like most natural habitats, the subsurface at the eFRC is a mosaic of spatiotemporally varying geochemical conditions that simultaneously apply distinct selection pressures on the same populations [3, 14, 15]. The population-level diversity preserved in the EnComs is therefore not merely residual noise but a structured reflection of these eco-evolutionary forces. Variants that are selectively neutral or mildly deleterious under one enrichment condition may confer a fitness advantage under another, providing a mechanistic basis for how environmental heterogeneity sustains genomic diversity at the population level [4, 37]. Crucially, the fact that the high-pN/pS signal of *napA* was consistent across enrichment conditions, rather than varying with the NO_3_^-^ gradient, suggests that the diversification of this gene is an enduring ecological signature of the population’s evolutionary history in a chronically NO_3_^-^-contaminated environment, rather than a transient response to the enrichment protocol.

### EnComs as a platform for dissecting eco-evolutionary dynamics across biological levels of organization

Enrichments along ecologically-relevant gradients and the systematic discovery of context-specific EnComs represents an approach that occupies a critical methodological niche between purely field-based environmental metagenomics and bottom-up synthetic community experiments. While the environmental metagenomics offers ecological realism but limited experimental control, experimentation with synthetic community offers control but sacrifices the ecological and genomic complexity of naturally-derived populations [33–35]. The ability to simultaneously interrogate community composition, interspecies correlation networks, metabolic potential, and population-level genomic evolution across replicated, defined chemical gradients provides a uniquely multiscale view of the eco-evolutionary forces shaping communities derived from a complex natural environment. Environmental metagenomics surveys have revealed the staggering breadth of microbial diversity across habitats [3, 23], but attributing specific functional roles to taxonomic variants remains deeply challenging in the absence of experimental control over the variables that shape fitness. Our approach partially addresses this interpretive gap by subjecting naturally-occurring ecotype diversity to defined selective pressures and observing the resulting community-and population-level responses.

However, our approach is not without limitations. The chemical gradients tested here, while calibrated to field-relevant ranges, are necessarily simplifications of the multiaxial and temporally dynamic geochemistry of the subsurface [15]. Additionally, although the enrichments in pH and nitrate gradients were replicated across different media (NLDM and SGW), the original inoculum was derived from a single homogenized sediment source. A few other processing and culturing steps may have introduced some bias in the composition of taxa in enrichments. For instance, spore-forming and dormancy-tolerant taxa may have better survived the freeze-thaw cycle during sediment processing and transport, and fast growers may have been favored by enrichments in nutrient replete media at 30°C, which is higher than *in situ* temperatures (∼14-18°C). Notwithstanding these limitations, our findings uncovered how environmental context dynamically shapes ecologically-relevant interactions among taxa that were consistent with previous findings, and substantiated by orthogonal evidence, e.g., MAG-encoded physiological capabilities of community members. Furthermore, the consistency between the taxa identified in the EnComs and those detected in amplicon-based surveys at the eFRC [26] supports the ecological relevance of the communities generated here, and the condition-specific community structure changes observed across gradients provide an experimental foundation for building and testing predictive models of subsurface biogeochemical cycling [15]. Future studies could improve and build on this framework by incorporating fluctuating chemical conditions to simulate temporal dynamics, extending enrichments to more transfers to probe longer-term evolutionary trajectories, and performing metatranscriptomics, metaproteomics, or metabolomics to characterize community-level physiology inferred from metagenome-encoded metabolic potential of constituent taxa.

Altogether, this work demonstrates that EnComs identified through high-throughput enrichments derived from a natural environmental source, characterized by long-read metagenomics across defined chemical gradients, can uncover the eco-evolutionary mechanisms, including enzymatic variants, metabolic complementarity, and condition-dependent selective pressures, that underpin niche partitioning and community assembly in complex microbial systems. These insights are foundational for developing predictive models of how natural microbial communities will respond to perturbations in nitrogen loading, pH, and carbon availability, conditions that are changing in subsurface environments globally as a result of agricultural contamination and climate-driven geochemical shifts [6, 7, 11].

## Materials and Methods

### Field sample collection

Subsurface sediment samples for enrichment were retrieved March 07, 2023 from the vadose zone of the SSO M6 well at a depth of 76-152 cm (30-60 inches; designated M6-C2) and sediment samples for amplicon sequencing were taken immediately below this from a depth of 152-229 cm (60-90 inches; designated M6-C3) (lat: N35-58-37.76/ long: W84-16-23.78) adjacent to the S-3 pond at the Oak Ridge Reservation ENIGMA Field Research Center (ORR eFRC) within the Y-12 National Security Complex in Oak Ridge, Tennessee, USA. Prior to borehole drilling, a 2.54 cm (1 inch) Geoprobe was advanced through the subsurface to collect a continuous core at 0.76 m (2.5 foot) intervals. Cores were capped, surveyed for radioactivity, placed in nitrogen atmosphere and stored at -80°C until enrichment. Soil analysis indicated a dry gravelly clay with 12.9% soil moisture and slight expansion to 85.1 cm (33.5 inches) recovery. NO_3_^-^ concentration and pH data from the eFRC were measured in a previous study by Smith et al. [14].

### Anaerobic plate enrichment

To investigate the effects of field-relevant stressors (e.g., pH, nitrate) and carbon availability on microbial assemblages, high-throughput (HT) enrichments were performed. Sediment samples (3g) from M6 were homogenized and mixed with 15mL of 30 mM phosphate buffer (amended with 5 mM pyrophosphate buffer) and sonicated for 30 minutes in a water bath to detach microbes. The resulting inoculum was then placed in an anaerobic glove box (Coy Lab Products, Ohio, USA) with an atmosphere composed of 75% N_2_, 20% CO_2_, and 5% H_2_ for 15 minutes to allow soil particles to settle before being added to each well. We utilized a minimal Synthetic Groundwater Medium (SGW) (details in **Supplementary Table 1**), and a defined, carbon-rich media, NLDM [38]. Synthetic groundwater was prepared according to a previous study [39], and the ingredients were as follows: FeSO_4_ (2 μM), MnCl_2_ (5 μM), Na_2_MoO_4_ (8 μM), MgSO_4_ (0.8 mM), NaNO_3_ (7.5 mM), KCl (0.4 mM), KNO_3_ (7.5 mM), CaCl_2_ (0.2 mM), NaH_2_PO_4_ (5 mM), and amended with vitamin mixtures at 10 mL/L.

The mineral mixture (pH 6.0) comprised 1.5 g L^-1^ NTA disodium salt, 3 g L^-1^ MgSO_4_ 7H_2_O, 0.5 g L^-1^ MnSO_4_ H_2_O, 1 g L^-1^ NaCl, 0.1 g L^-1^ FeSO_4_ 7H_2_O, 0.1 g L^-1^ CaCl_2_ 2H_2_O, 0.1 g L^-1^ CoCl_2_ 6H_2_O, 0.13 g L^-1^ ZnCl, 0.01 g L^-1^ CuSO_4_ 5H_2_O, 0.01 g L^-1^ AlK(SO_4_)2 12H_2_O, 0.01 g L^-1^ AlK(SO_4_)2 12H_2_O, 0.01 g L^-1^ Boric Acid, 0.025 g L^-1^ Na_2_MoO_4_ 2H2O, 0.024 g L^-1^ NiCl_2_ 6H_2_O, 0.025 g L^-1^ Na_2_WO_4_ 2H_2_O, and 0.02 g L^-1^ Na_2_SeO_4_. The vitamin mixture comprised 2 mg L^-1^ of d-biotin, 2 mg L^-1^ folic acid 10 mg L^-1^ pyridoxine HCl, 5 mg L^-1^ riboflavin, 5 mg L^-1^ thiamine, 5 mg L^-1^ nicotinic acid, 5 mg L^-1^ pantothenic acid, 0.1 mg L^-1^ vitamin B12, 5 mg L^-1^ p-amino benzoic acid, and 5 mg L^-1^ alpha-lipoic acid.

For SGW plates, multifactorial combinations of NO_3_^-^ (0-40 mM) as an electron acceptor and carbon (20 mM; glucose, pyruvate, lactate, citrate, and acetate) as an electron donor were arrayed. All NO_3_^-^ (1 M) and carbon stocks (2 M) were prepared anaerobically. Three different pH values (4.5, 6, and 7), reflecting the field range, were also incorporated. For NLDM plates, which already contain diverse carbon sources, no additional carbon was added. Instead, a multifactorial combination of pH (4.5, 6, and 7) and NO_3_^-^ (0-40 mM) was tested. Detailed plate layouts for the HT enrichments are provided in **Supplementary Table 1**.

Each well was inoculated with 40µL of M6 inoculum and 360µL of corresponding media mix. Plates were incubated without shaking at 30°C for 7 days in an anaerobic glove box (Coy Lab Products, Ohio, USA) with an atmosphere composed of 75% N_2_, 20% CO_2_, and 5% H_2_. After 7 days, 40µL was transferred from each well into a new plate with the exact same conditions to promote continued microbial growth. Plates were then centrifuged at 3600 rpm for 10 minutes to form cell pellets, and the remaining liquid was discarded. The plates were stored at -80°C until sequencing. A total of three transfers were performed over four weeks. Glycerol stocks were prepared from the last transfer using 50% glycerol and 50% base medium and stored at -80°C for future analysis.

### DNA extraction and long-read sequencing

To extract DNA from the enrichments, plates were thawed and each sample was extracted using the Zymo *Quick*-DNA HMW MagBead Kit. DNA concentrations were quantified using a Qubit fluorometer. Sequencing libraries were prepared using the Nanopore Rapid Barcoding Kit 96 V14 (Oxford Nanopore Technologies, Oxford, UK). The first batch of samples (n=96, SGW T3) were sequenced using the MinION Mk1C (Oxford Nanopore Technologies, Oxford, UK). The remaining samples were sequenced using the PromethION P2 Solo (Oxford Nanopore Technologies, Oxford, UK) to achieve deeper coverage. Raw pod5 files from the Nanopore sequencing were basecalled and demultiplexed using Dorado (Oxford Nanopore Technologies, Oxford, UK) with the super high accuracy basecalling model dna_r10.4.1_e8.2_400bps_sup@v5.0.0. Reads were trimmed to remove low quality reads and ends using BBDuk [81] with the settings qtrim=rl, trimq=10, and maq=10. All metagenomic sequencing data generated in this study are publicly available through https://narrative.kbase.us/narrative/262281. DNA was extracted from sediment cores and amplicons were sequenced for ASV analysis as previously described by Goff et al. [26] Amplicon sequence data is publicly available in KBase narratives (https://kbase.us/n/112150/61/ and https://narrative.kbase.us/narrative/262281).

### Metagenomic assembly, mapping, and profiling

Reads from each sample were grouped based on the enrichment medium (NLDM or SGW) and concatenated for co-assembly. The assemblies were generated using metaFlye [82] for each subset of samples. A combined assembly of both NLDM and SGW samples was generated by combining the two assemblies using metaFlye in subassembly mode. Each individual sample was then mapped to the co-assembly using BBMap [81] in long-read mode (mapPacBio.sh) with standard settings. The data was aggregated using Anvi’o (version 8, last updated and verified with version 9) [41, 83] by generating a contigs database from the assembly (anvi-gen-contigs-database) and generating individual profiles for each sample from the mapping data (anvi-profile). During sample profiling, single nucleotide variants (SNVs), single amino acid variants (SAAVs), and single codon variants (SCVs) were identified for each base and codon position in the assembly. All of the profiles were merged to generate a singular metagenomic profile database (anvi-merge). Metagenomics databases, including gene calls, coverage, and variants that were generated using Anvi’o, and summary data is available at https://github.com/baliga-lab/sediment-encoms.

### Metagenomic annotation, binning, and refinement

Gene calling was performed during contigs database generation with Anvi’o [41] using Prodigal [84]. Additionally, a Hidden Markov Model was run to identify marker genes (anvi-run-hmms) [85], and taxonomy was estimated for each marker gene (anvi-run-scg-taxonomy) [86]. Each coding sequence was annotated with NCBI COGs (anvi-run-ncbi-cogs) [54] and KEGG Kofams (anvi-run-kegg-kofams) [55]. Binning was done manually in Anvi’o’s interactive mode (anvi-interactive) following the clustering generated during profile merging that orders contigs based on similarity of GC content and read coverage across samples. All genes in the assembly were binned, with attention paid to maximize completeness while minimizing contamination for bins that appear to represent whole genomes [41]. Completeness and contamination of each MAG was verified with CheckM [87] implemented in Kbase [88], and each bin is available for analysis in https://narrative.kbase.us/narrative/262281. Metabolic pathway completeness assessment was conducted in Anvi’o (anvi-estimate-metabolism) [89]. The Anvi’o databases containing the merged, binned, and annotated metagenomes are available along with a summary of the annotations and coverages for each bin/MAG. The relative abundance of each bin was calculated using the Q2Q3 (inner quartiles, to reduce noise from outliers) mean coverage of a bin in each sample as a fraction of the total coverage of all bins. The analyses and visualizations were conducted using custom scripts in Python v3.14.3 available along with underlying data at https://github.com/baliga-lab/sediment-encoms.

### Statistical analyses

Statistical analyses were performed using Pandas [90], NumPy [91], SciPy [92], and NetworkX^93^. Visualizations were generated using Matplotlib [94], Seaborn [95], and NetworkX [93]. Two-sided independent t-tests were performed using scipy.stats.ttest_ind, and Welch’s t-tests were performed when appropriate by setting equal_var to False. Two-sided linear least-squares regressions were calculated using scipy.stats.linregress. Significance was determined if tests returned p-values less than 0.05. Additional tools used for exploratory analyses include Scikit-learn [96] and Scikit-bio [97]. All analyses were conducted in Python-based Jupyter Notebooks (https://github.com/baliga-lab/sediment-encoms).

### Correlation network analysis

We performed linear regression analysis in Python using SciPy [92] to inspect the correlation of relative abundance of bins with NO_3_^-^ concentrations and pH. We performed T-tests to evaluate differences in abundance of genera between conditions using SciPy (ttest_ind). We generated a matrix of the mean abundance of each bin in each sample to estimate co-occurrence using Fastspar [46] which is an optimized implementation of the SparCC algorithm [45]. To calculate the significance of the correlation between each bin, we implemented bootstrapping (1000 iterations of randomly shuffled real data) following the Fastspar documentation (https://github.com/scwatts/fastspar) to calculate exact p-values for each pair of bins. We used this approach to identify significant correlations between bins across all conditions as well as in mindful conditional subsets. We ran each subset of conditions individually and applied bootstrapping each time. We analyzed the networks of significant correlations using custom Python-based Jupyter Notebooks which used Pandas [90], Numpy [91], SciPy [92], and NetworkX [93]. Link-community analyses were conducted using the bootstrapped correlation data generated by Fastspar using the python script link_clustering.py written by Ahn et al. [47] (https://github.com/bagrow/linkcomm/blob/master/python/link_clustering.py). Jupyter Notebooks containing all analyses are openly available at https://github.com/baliga-lab/sediment-encoms.

### MAG metabolic comparisons

Metabolic pathway completeness estimations for each MAG were conducted using Anvi’o (anvi-estimate-metabolism) [89]. The output was analyzed using a custom Python-based Jupyter Notebook (https://github.com/baliga-lab/sediment-encoms). For each MAG, the stepwise pathway completion percentage was used to determine where a pathway was complete (100%) or partial (<100%). Percentage metabolic “overlap” was calculated pairwise as the intersection of pathways p encoded by both MAGs i and j divided by the number of pathways encoded by one of the MAGs n_i_:

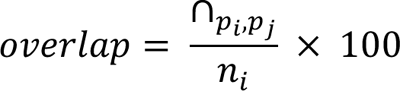

The percentage “similarity” of the MAGs was calculated as the intersection of pathways encoded by both MAGs divided by the union of all pathways encoded by both MAGs:

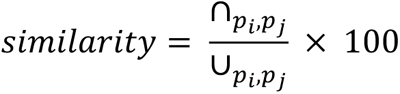

These calculations were done either for complete pathways only or for the total number of all complete and partial pathways.

### Population genetic variability (pN/pS) analysis

We exported data tables including SCVs, pN, and pS for each codon position for the Bin 1 *Neorhizobium* in each sample for each condition from Anvi’o (anvi-gen-variability-profile) [37]. We calculated pN/pS for each codon position using the consensus pN and pS values from genes that had a coverage of at least 10 and a minimum departure from consensus of 0.1. We generated mean pN/pS values for each gene using the Anvi’o script “anvi-get-pn-ps-ratio” which implements a normalization when averaging based on potential mutations for each codon [37]. We used the gene-level data to calculate the genome-wide average values and the codon-level data for position-specific pN/pS values for each sample. We calculated statistical differences between conditions using SciPy (ttest_ind and linregress) [92] and visualized the data in a Python-based Jupyter Notebook (https://github.com/baliga-lab/sediment-encoms).

## Acknowledgements

This material by ENIGMA-Ecosystems and Networks Integrated with Genes and Molecular Assemblies (http://enigma.lbl.gov), a Science Focus Area Program at Lawrence Berkeley National Laboratory is based upon work supported by the U.S. Department of Energy, Office of Science, Office of Biological & Environmental Research under contract number DE-AC02-05CH11231.

